# Starvation drives co-existence in cross-feeding bacterial populations

**DOI:** 10.1101/2025.04.22.650053

**Authors:** William F. McLaughlin, Megan G. Behringer

## Abstract

Cross-feeding is a universal feature of natural microbial communities that extends even into subpopulation structures, generating cooperative ecotypes. However, we still lack an understanding of the ecological parameters that support the ecotype co-existence necessary for co-evolution between partners within populations. Here, we show that repeated moderate starvation – conditions that mimic the fluctuations experienced by microbes in nature – can drive frequency-dependent co-existence for hundreds of generations, providing a new mechanism for the origin of cooperation and the maintenance of population-level diversity in natural microbial communities.

## MAIN

Cross-feeding, or the exchange of resources between genotypically or phenotypically distinct individuals (Fritts et al., 2021), is a ubiquitous form of cooperation within natural microbial communities (Kost et al., 2023), ranging from interdomain to intraspecific interactions. Bacterial populations often display cross-feeding through metabolic partitioning, with subpopulations, or ecotypes, that have evolved to specialize on certain resources or produce certain products utilized by others in a community (Blount et al., 2012; Finkel & Kolter, 1999; Helling et al., 1987; Rainey & Travisano, 1998). The initial step towards cross-feeding is proposed to be an ecotype developing a facultative or obligate auxotrophy for some resource provided by prototrophic individuals in their environment (Morris et al., 2012; D’Souza & Kost, 2016). Prototrophic producers can grow independently but bear the metabolic burden of supporting the population, whereas auxotrophs expend no energy making a certain resource, but can grow only when that resource is provided exogenously. This tradeoff associated with these lifestyles can vary greatly between resources and environments, leaving the parameters that support the long-term co-existence of auxotrophs and producers that comprise microbial networks poorly understood.

Previous work has shown that altered resource availability can modulate the fitness and stability of obligate and facultative cross-feeding, but has assumed a constant, albeit limited, availability of resources through rapid short-term transfers or in chemostat cultures (Hoek et al., 2016; Sexton & Schuster, 2017). However, the intermediate disturbance hypothesis (Connell, 1978; Grime, 1973) posits that diversity peaks when turnover occurs at a moderate frequency—neither too often nor too rarely. This suggests that longer periods between transfers or resource replenishment could influence the success of different subpopulations, a phenomenon observed in microbial communities (Kassen et al., 2000) that has recently been predicted to drive dynamic co-existence patterns in multispecies microbial systems (Narla et al., 2025). Additionally, moderate repeated cycles of feast and famine can support the emergence of facilitative cross-feeding ecotypes that co-exist for hundreds of generations (Behringer et al., 2018, 2022). By subjecting cross-feeding populations to feast-famine cycles of different lengths, we can investigate how one-way feeding stability is shaped by the complex stressors associated with resource fluctuations that are experienced in nature. We hypothesized that increasing the periodicity between resource replenishment will balance lifestyle trade-offs associated with metabolic partitioning, stabilizing obligate cross-feeding on longer timescales.

To test this, we engineered one-way methionine cross-feeding populations of *Escherichia coli* K-12. This system consisted of mutant strains auxotrophic Δ*metB*, which lacks the first gene in the methionine *de novo* biosynthesis pathway (Duchange et al., 1983; Tran et al., 1983), and overproducer Δ*metJ*, which lacks the repressor of *de novo* methionine biosynthesis (Saint-Girons et al., 1984; Smith et al., 1985; Smith & Greene, 1984) (**Fig. 1A**). To verify that the co-culture system enables the growth of the auxotroph only when cultured with a complementary overproducer, each strain was grown in modified M9 minimal medium (mM9) conditioned by the wild-type or overproducer Δ*metJ* strain (filtered spent media following 24 hours of growth and re-supplemented with 0.2% glucose). The wild-type conditioned medium only supports the growth of the Δ*metJ* and wild-type strains, whereas the overproducer Δ*metJ* conditioned medium supports all three strains, rescuing auxotroph Δ*metB* growth (**Fig. 1B**).

**Figure 1.**
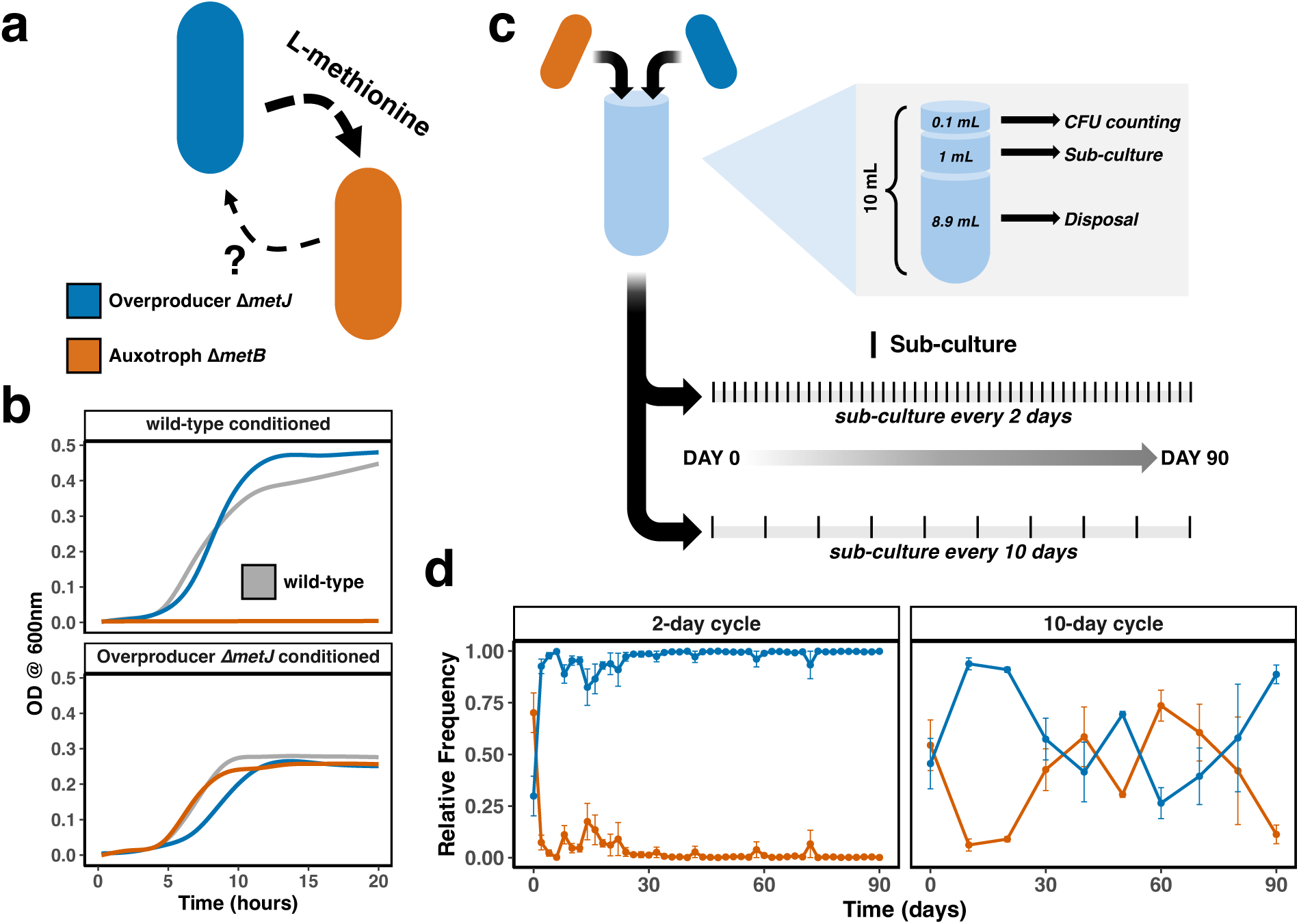
Engineered cross-feeder design and co-culture experiment. (A) Schematic of engineered mutant strains of *E. coli* K-12 composed of a methionine overproducer Δ*metJ* (blue) and a methionine auxotroph Δ*metB* (orange). Bold dotted arrow indicates the obligate exchange of methionine from overproducer to auxotroph; a small dotted arrow with a question mark denotes possible reciprocal externalization of resources. (B) Growth curves (measured by optical density at a wavelength of 600 nm) for both mutant strains and wild-type in the conditioned media of the overproducer Δ*metJ* and wild-type. Gray shading indicates 95% confidence intervals. (C) Experimental design and sampling schedule for 90 days of culture in starved and un-starved conditions. See Methods for culture conditions. (D) Mean relative frequency of each ecotype across 90 days of co-culture. Error bars indicate (+/-) SEM.

Next, we cultivated 12 replicate cross-feeding populations consisting of the auxotroph Δ*metB* and overproducer Δ*metJ* strains co-cultured in mM9 minimal medium (0.2% glucose) for 90 days and subjected them to a 1:10 bottleneck into fresh media either every 2 or 10 days (**Fig. 1C**). Using mean colony forming units (CFUs) per mL across populations 1, 2, and 3 (**Fig. S1)**, we tracked the relative frequency of each ecotype as the percentage of overall cell density for the duration of the experiment. Populations cultivated under 10-day transfer conditions exhibited divergent oscillations in ecotype frequencies (**Fig. S2**), suggesting negative frequency-dependent long-term co-existence (**Fig. 1D**). Alternatively in the 2-day transfer condition, the overproducer Δ*metJ* outperforms its auxotrophic counterpart (**Fig. S2**), converging upon equilibrium frequencies where the overproducer comprises the majority of the population (**Fig. 1D**). We calculated the steady-state ecotype frequency equilibrium (frequency where selection rate = 0) for both conditions by fitting a linear regression model to each ecotype’s selection rates across the final 60 days of co-culture after co-existence patterns manifest. In 2-day cycles, we predict ecotype equilibrium frequencies of 99% overproducer Δ*metJ* and 1% auxotroph Δ*metB* (**Fig. S3**). Alternatively, in 10-day cycles, we predict ecotype equilibrium frequencies of 56% overproducer Δ*metJ* and 44% auxotroph Δ*metB* (**Fig. S3**). However, because 10-day conditions promote steep negative frequency-dependent fitness (**Fig. S3**), the ecotypes ultimately oscillate around these frequencies, never converging upon an equilibrium (**Fig. S2**).

We quantified the costs and benefits associated with the lifestyles of each mutant over 10 days to discern why equilibrium frequencies and ecotype dynamics differ so greatly between the 2-day and 10-day culture conditions. We assessed the benefit gained by the auxotroph Δ*metB* not synthesizing methionine when it is exogenously supplied by competing it against the wild type in mM9 + 5.9 mM methionine (concentration based on Sezonov et al., 2007). Here, the auxotroph Δ*metB* strain is initially outperformed by the wild-type (**Fig. 2B**), before exhibiting an increasingly positive selection rate as resource limitation intensifies over 10 days (**Fig. 2C**). We then assessed the cost of overproducing methionine incurred by the overproducer Δ*metJ* in unsupplemented medium where released excess methionine would provide a communal benefit. In these conditions, the overproducer Δ*metJ* performs poorly relative to wild-type (**Fig. 2B**) and exhibits an increasingly negative selection rate as resource limitation intensifies (**Fig. 2C**).

**Figure 2.**
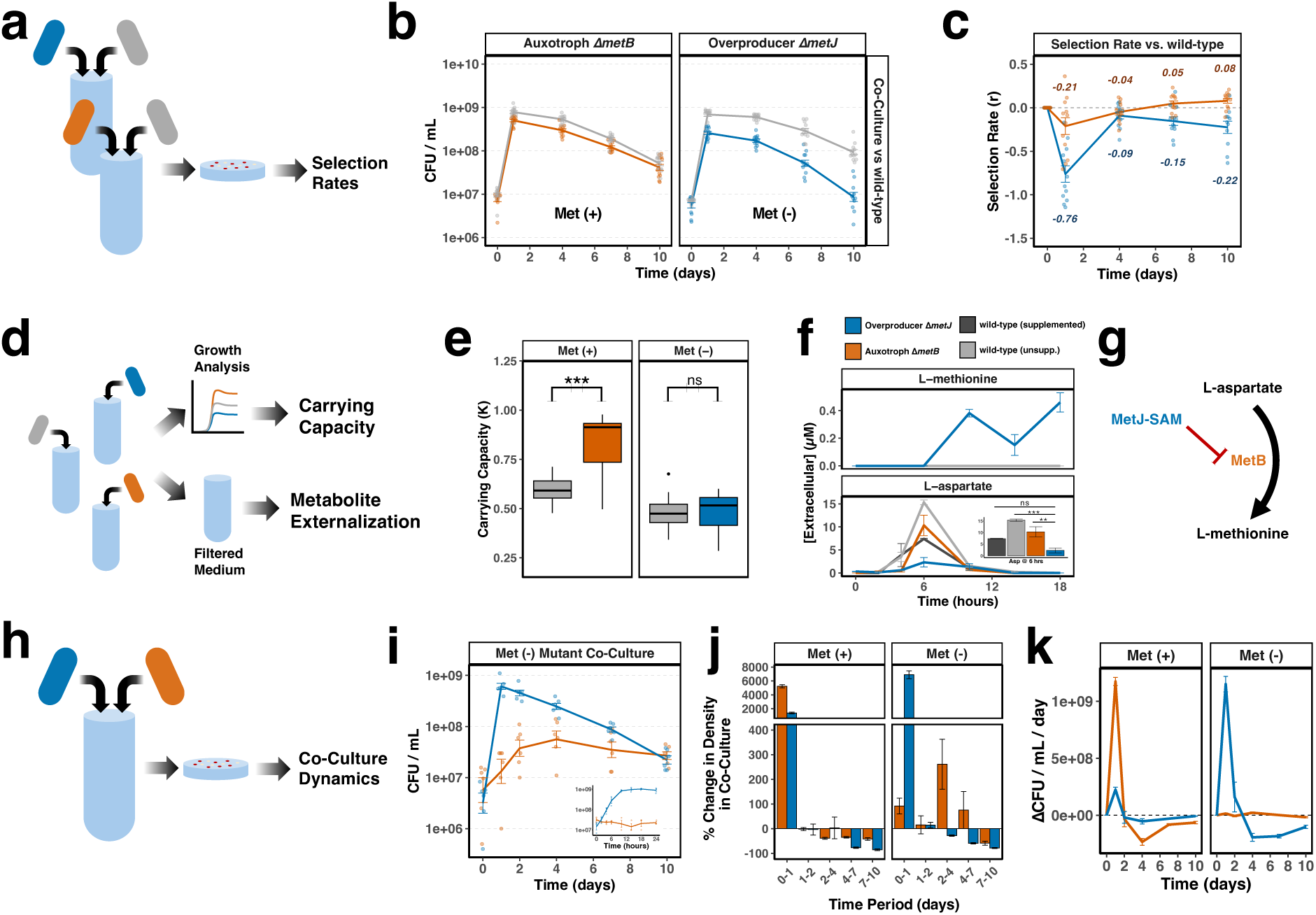
Cross-feeding system parameterization. (A) Schematic of selection rate co-culture experiments. (B) CFU/mL between each mutant and wild-type (n=12). Error bars indicate (+/-) SEM. (C) Change in selection rate (*r*) between clones of either overproducer Δ*metJ* (blue) in mM9 or auxotroph Δ*metB* (orange) in mM9+methionine cultured with a wild-type background across 10 days (n=12). Error bars indicate (+/-) SEM. (D) Schematic of growth analysis and metabolite externalization experiments. (E) Carrying capacity of each mutant and wild-type in mM9 +/- 5.9 µM L-methionine (n=18). Bars indicate 95% confidence intervals. (F) LC-MS quantification of extracellular amino acids (n=3 for each mutant, n=6 at hour 18). See Methods for sample preparation and analysis. Figure-in-figure indicates aspartate levels at six hours. Error bars indicate (+/-) SEM. (G) Graphic of methionine biosynthesis and protein regulation. (H) Schematic of mutant co-culture experiments. (I) CFU abundance of clones (n=6) of overproducer Δ*metJ* or auxotroph Δ*metB* co-cultured directly across 10 days. Figure-in-figure indicates growth across a 24-hour period. Error bars indicate (+/-) SEM. (J) Density change represented in percent abundance per day in co-culture in mM9 medium +/- methionine (n=9). Error bars indicate (+/-) SEM. (K) Rate of CFU change per day in co-culture in mM9 medium +/- methionine (n=9). Error bars indicate (+/-) SEM.

To further characterize the phenotypes underlying the lifestyle tradeoffs in these conditions, we measured the carrying capacity and externalization of L-methionine and its precursor, L-aspartate, in each mutant and wild-type over 18 hours (**Fig. 2D**). In unsupplemented media, the overproducer Δ*metJ* exhibits a carrying capacity comparable to the wild-type. Alternatively, when supplementing methionine, the auxotroph Δ*metB* exhibits a significant improvement in carrying capacity over the wild-type (Welch’s *t* = 5.3357, *P* = 2.075e-05) (**Fig. 2E**). This improvement in carrying capacity can be further explained by metabolite externalization.

Specifically, concentrations of the methionine precursor L-aspartate spike in all cultures at 6 hours before dropping to undetectable levels by 18 hours (**Fig. 2F**), likely due to re-internalization and utilization as a carbon source or for amino acid biosynthesis. These changes correspond with inflections in the growth curve of methionine supplemented cultures as glucose is depleted at 6 hours, triggering a diauxic shift and eventual transition to stationary phase around 12 hours (**Fig. S4**). Since the auxotroph Δ*metB* is unable to convert L-aspartate to L-methionine, re-internalized aspartate may be redirected towards biomass production, contributing to its increased carrying capacity (**Fig. 2E**). Alternatively, L-aspartate externalization is significantly lower in the overproducer Δ*metJ* compared to unsupplemented wild-type and auxotroph Δ*metB* at 6 hours (ANOVA, Tukey’s HSD both *P* < 0.01) (**Fig. 2F**), as it is likely being funneled into L-methionine production due to the lack of negative regulation of methionine *de novo* biosynthesis by MetJ protein (Liu et al., 2015) (**Fig. 2G**). As a result, the overproducer Δ*metJ* is the only strain to release appreciable amounts of L-methionine by 18 hours as cells enter stationary phase (**Fig. 2F**), albeit at a lower cell density (**Fig. S4**). Together, these results highlight the tradeoff: overproducer Δ*metJ* can grow autonomously but incurs a metabolic cost from constitutive methionine production, while auxotroph Δ*metB* requires external methionine to survive but benefits from avoiding this biosynthetic burden, resulting in higher biomass yield when supplemented.

To assess how lifestyle tradeoffs in methionine production manifest when the mutant strains are simultaneously present, we quantified the abundances of each ecotype over 10 days of co-culture in mM9 medium without supplemented methionine (**Fig. 2H**). The overproducer Δ*metJ* achieves its peak density at day 1 but steadily declines thereafter over the 10-day culture. Alternatively, after one day, the auxotroph Δ*metB* comprises only ∼1% of the population (**Fig. 2I, Fig. S5**), but continues to increase in abundance starting on day 2 before peaking at day 4 and sustaining its abundance for the duration of the 10-day culture. This suggests that the availability of methionine may not reach critical levels for auxotroph Δ*metB* growth until cells reach late-stationary phase and begin to enter death phase. Ultimately, by day 10, the overproducer Δ*metJ* and auxotroph Δ*metB* exhibit similar abundances (Welch’s *t* = 0.7432, *P* = 0.4745) (**Fig. 2I**), accounting for the stark differences in ecotype dynamics between 2-day and 10-day transfer cycles.

To confirm the effect of overproducer Δ*metJ* death on the delayed success of auxotroph Δ*metB*, we removed auxotroph Δ*metB* dependence on overproducer Δ*metJ* by co-culturing the mutants for 10 days in mM9 medium with supplemented methionine and determining each ecotype’s density change between each sampling period. When methionine is supplemented, the auxotroph Δ*metB*’s growth and survival more closely resemble the overproducer Δ*metJ* (**Fig. 2J, Fig. S5**). As such, one factor contributing to the increased long-term survival of the auxotroph Δ*metB* may be its more conservative growth rate when overproducer Δ*metJ* acts as the sole source of methionine (**Fig. 2K**), a pattern consistent with a balance between growth rate and cellular maintenance in *E. coli* (Biselli et al., 2020). Taken together, this suggests the initial rapid growth and death of the overproducer Δ*metJ* supports the co-existence of the auxotroph Δ*metB* by facilitating a slower auxotroph Δ*metB* growth rate, helping to maintain auxotroph Δ*metB* density during starvation.

The effect of metabolic similarity (or dissimilarity) is well understood in microbial ecologies, with divergent niches and complementary metabolisms yielding the greatest co-existence (Chuang et al., 2024; Giri et al., 2021; Harcombe et al., 2018). Yet the metabolisms of related organisms are highly congruent, meaning that cross-feeding within populations must overcome intraspecific competition to persist. We demonstrate the role dynamic systems play in maintaining intraspecific diversity, with temporal changes in resource availability playing an additional part in promoting ecotype co-existence. Rapid growth and limited methionine release by the overproducer depletes the resource pool and slows the growth of its auxotrophic partner, allowing the overproducer to dominate in short transfer regimes. Later succession of available nutrients as cells die and lyse eliminates this effect, allowing for the equal abundance of ecotypes to be achieved at moderate intensities of starvation around 7-10 days of culture. This observation provides support for the intermediate disturbance hypothesis (Connell, 1978; Grime, 1973), with increased refeeding disturbance periodicity balancing the abundance of ecotypes of different competitive capacity. We also provide empirical evidence for how subpopulations have been predicted to favor dividing labor into rapid reproducers and slow-growing secondary metabolite specialists (Kumakura et al., 2023), highlighting the importance of gleaner-opportunist trade-offs (Fredrickson & Stephanopoulos, 1981; Grover, 1991; Yamamichi & Letten, 2022) and relative nonlinearity in growth capacity (Chevin et al., 2022) in the maintenance of population-level diversity in bacteria. These ideas were developed in multispecies microbial systems (Fredrickson & Stephanopoulos, 1981), but diversity does not stop at the species level, and we show that the eco-physiological feedback induced by one subpopulation’s death as resources grow scarce can prime another’s slow-growth and survival. This phenomenon creates storage effects that reveal themselves in temporal patches, where the auxotrophic lifestyle can thrive in later patches by being forced to display conservative growth in early ones.

Most strikingly, increased refeeding periodicity can promote dynamic ecotype co-existence and maintain obligate intraspecific cross-feeding for extended periods. The pattern observed in the 10-day cycle co-culture indicates negative frequency dependence (NFD) and aligns with predictions that strong NFD will drive divergent oscillations (Chevin et al., 2022). In 2-day cycles, the high density in the overproducer Δ*metJ* suppresses invasion by the auxotroph Δ*metB* and creates a priority effect driven positive frequency-dependence (Ke & Letten, 2018). In all, refeeding periodicity and starvation intensity can drive two alternative states in the same system. With natural microbial environments regularly experiencing widely varied levels of stress or disturbance periodicity, we provide a new paradigm for understanding how the nascent stages of mutualism are stabilized, and how cooperation may emerge in natural populations of bacteria.

## Materials & Methods

### Strain construction

The genes of interest for construction of the cross-feeding strains were identified from a search of genes annotated as essential for growth in minimal medium via the EcoCyc database (Moore et al., 2024). PFM2, a prototrophic derivative of the *E. coli* K-12 str. MG1655 reference strain (Lee et al., 2012), served as the wild-type background for all constructed strains. Using *E. coli* K-12 deletion strains from the Keio Collection (Baba et al., 2006), the Δ*metB*726::*kan* mutation was introduced into PFM2 via P1 transduction (Saragliadis et al., 2018) using *E. coli* K-12 BW25113::Δ*metB* as the donor to create the auxotroph Δ*metB* strain. The Δ*metJ*725::kan mutation was transduced from *E. coli* K-12 BW25113:: Δ*metJ* to create the overproducer Δ*metJ* strain. This was accomplished by expressing the FLP recombinase from the helper plasmid pCP20, which resulted in the removal of the kanamycin resistance cassette leaving behind the FRT scar site (Datsenko & Wanner, 2000). Kanamycin resistance was left in the auxotrophic Δ*metB* strain in order to count cells below detectable limits when plated in co-culture. The overproducer Δ*metJ* strain also carries a Δ(araD-araB)567 mutation that renders the strain *araBAD*(-), while the auxotroph Δ*metB* strain is *araBAD*(+). Here, the *araBAD l*ocus acts as a neutral marker, allowing the Δ*metJ and ΔmetB* strain to be discernible from each other on tetrazolium arabinose (TA) agar (10 g/l tryptone, 1 g/l yeast extract, 5 g/l NaCl, 16 g/l agar, 10 g/l L-arabinose, 0.005% tetrazolium chloride) as red and white colonies, respectively.

### Growth curves

Strains to be measured as a growth curve were grown overnight in 2 mL of modified M9 minimal medium (mM9) (119.39 mM disodium phosphate (anhydrous), 55.11 mM monopotassium phosphate, 21.39 mM sodium chloride, 46.74 mM ammonium chloride, 1 mM magnesium sulfate, 0.1 mM calcium chloride, and 20% w/v glucose) shaking at 180 rpm at 37 °C in 16 x 100 mm borosilicate glass culture tubes. Auxotroph Δ*metB* strains were supplemented with 5.9 mM L-methionine in all preparatory overnights, according to the concentration of L-methionine in LB broth (Sezonov et al., 2007). 1 mL of culture was then spun down in a microcentrifuge for 60 seconds at 10,000 rpm. Supernatant was decanted, and the pellets were resuspended in 1mL of fresh mM9 minimal media. The resuspended cells were re-spun for 60 seconds at 10,000 rpm, decanted, and again resuspended in 1mL of mM9 minimal media. These washing steps were necessary to remove all excess waste products and supplemented methionine from the overnights.

Washed overnight cultures were then inoculated with a 1:10 dilution into fresh mM9 media to recreate the transfer volume during the 90 day co-culture experiment. Diluted cultures were aliquoted 100 uL each into wells of a Greiner Cellstar 96-well cell culture plate (Product #: 655180) and placed into a plater reader for measurement. To generate conditioned media, strains used to condition the media were inoculated from a single colony in 500 mL of fresh mM9 minimal media and grown overnight for 24 hours shaking at 180 rpm at 37 °C in a 1L Erlenmeyer flask. The media was then filtered using Fisherbrand 500 mL filter unit (0.2 µm aPES membrane) (Product #: FB12566504). For growth curves in spent media, cultures were grown overnight for 18 hours shaking at 180 rpm at 37 °C in 2 mL of spent media that had been resupplemented with 0.2% glucose. These cultures were subjected to the same wash steps as before with fresh mM9. The washed cultures were then inoculated with a 1:100 dilution into filtered conditioned media and 100 µL of the diluted culture aliquoted into each well of a Greiner Cellstar 96-well cell culture plate (Product #: 655180) and placed into a plate reader for measurement. All growth curves were measured using Agilent Biotech Epoch 2 plate reader at 180 rpm double orbital shaking at 37 °C for 20 hours, recording absorbance readings at 600 nm every 15 min. Growth curve parameters (carrying capacity, growth rate, etc.) were quantified using the R package *growthcurver* version 0.3.1 (Sprouffske & Wagner, 2016).

### Culture conditions, co-culture experiment, and pairwise fitness experiments

An ancestor glycerol stock was streaked onto LB-Miller agar (10 g/l tryptone, 5 g/l yeast extract, 10 g/l NaCl) for each mutant strain. Experimental starting cultures for each culture replicate were established as follows: For each strain, 12 colonies were selected and inoculated into individual 16 x 100 mm glass culture tubes containing 10 mL of LB-Miller broth and grown overnight for 18 hours shaking at 180 rpm at 37 °C. To preserve a copy of the experimental starting cultures, a glycerol stock was prepared for each of the 12 overnight cultures for each mutant (200 µL 70% glycerol, 1 mL culture). To initiate the experimental co-cultures, 1 mL of culture was then spun down in a microcentrifuge for 60 seconds at 10,000 rpm. Leftover media was decanted, and the pellets were resuspended in 1mL of fresh mM9 minimal media. The resuspended cells were re-spun, decanted, and again resuspended in 1mL mM9 minimal media. These washing steps were necessary to remove all excess waste products and leftover amino acids in LB from the overnights to have the co-culture experiment start with fresh, consistent conditions. Resuspended cells were adjusted to an OD of 0.4 at 600 nm across all cultures. Each of the 12 co-culture replicates were inoculated with 100 µL of the corresponding ancestor 1-12 for both mutants for each replicate.

For the remainder of the 90 day co-culture experiment, the cultures were grown in aerobic conditions in mM9 minimal media supplemented with 0.2% glucose in 16 x 100 mm glass tubes shaking at 180 rpm at 37 °C. At each time point (either every 2- or 10-days) the tube was homogenized by pipetting to mix, and a subculture of 1 mL was inoculated into 9 mL of fresh mM9 media. Two cell pellets and a glycerol stock were collected and frozen every transfer for the first 30 days of co-culture, then every 10 days through day 90. Strains from the co-culture are stored at −80° C and are made available upon request. CFU data were collected by spot-plating 5 µL of diluted co-culture onto both tetrazolium agar and LB + 50 mg/mL kanamycin. For any single plating timepoint where cells were uncountable during the 90 day continuous culture experiment, a count of 0.1 was input at the same dilution factor as other replicates for that particular ecotype.

We utilized competitive fitness experiments to quantify selection rate (*r*) (Lenski et al., 1991; Levin et al., 1977). For pairwise co-culture dynamics and competitive fitness experiments, strains were grown overnight in 10 mL of mM9 liquid media shaking at 180 rpm for 18 hrs at 37 °C in 16 x 100 mm borosilicate glass culture tubes. 1 mL of culture was spun down in a microcentrifuge for 60 seconds at 10,000 rpm. Supernatant was decanted, and the pellets were resuspended in 1mL of fresh mM9 minimal media. The resuspended cells were re-spun, decanted, and again resuspended in 1mL of mM9 minimal media and adjusted to an OD of 0.4 at 600 nm across all cultures. To each tube, 100 µL of Δ*metB* and 150 µL Δ*metJ* were added to each tube and plated for CFU at time zero (Δ*metJ* mutants had an observable higher mortality rate in mM9, therefore starting volumes were adjusted to achieve an equal proportion of viable cells at the initial time point). For competitions that required supplemented methionine, 5.9 mM methionine was added to the media (concentration determined from HPLC of enriched LB media (Sezonov et al., 2007)). Cells were plated on tetrazolium agar (TA) to differentiate cells for colony forming units counting on days 0, 1, 2, 4, 7, and 10. Plating of culture on TA plus arabinose agar turns araBAD+ colonies pink and araBAD− colonies red for easy enumeration of genotypes. Selection rate (*r*) was determined as the difference of the natural log of the ratio of each competitor’s CFU counts at the final and initial time points:

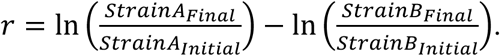

### Metabolomic quantification of amino acids

Strains to be measured were grown overnight in 10 mL of mM9 liquid media shaking at 180 rpm for 24 hours at 37 °C in 16 x 100 mm glass culture tubes (Δ*metB* was grown with 5.9 mM methionine supplement). Without washing, 1 mL of three biological replicates of Δ*metJ*, Δ*metB*, and wild-type cultures were inoculated in 9 mL of mM9 minimal media in a glass tube either supplemented with 5.9 mM methionine (Δ*metB*, wt) or unsupplemented (Δ*metJ*, wt) and resampled from the same tube throughout the growth period. 0.5 mL samples were collected at hours 0, 2, 4, 6, 10, 14, and 18 based on growth curves following a 1:10 dilution from an 18 hour culture into fresh methionine-supplemented or unsupplemented mM9 minimal liquid medium (**Fig. S4**). 0.5 mL of culture was spun through Amicon Ultra 0.5 mL Ultracel 10K centrifugal filters (Product #: UFC501096) in a microcentrifuge at 10,000 rpm for 5 minutes to remove biological material. Samples were flash frozen in liquid nitrogen and stored at −80 °C. Samples were then submitted to the Vanderbilt Mass Spectrometry Research Center at Vanderbilt University for LC-MS metabolomics quantification. Values below or above the lower limit of calibration (0.2 µM < concentration < 2 mM) were filtered out.

### Data Availability

All code for data analysis and production of figures are deposited on GitHub at https://github.com/BehringerLab/Ecotype_Coexistence.

## Acknowledgments

We would like to thank J. McKinlay, N. Komarova, and K. Catania for their helpful comments and suggestions leading up to the publication of this study. Metabolomic sample prep and quantification was performed at the Vanderbilt Mass Spectrometry Research Center (MSRC). This work was supported by National Institute of General Medical Sciences Grant R35GM150615 (M.G.B.), as well as additional funds provided by the Evolutionary Studies Initiative at Vanderbilt (W.F.M. and M.G.B).

## Supplementary Figures

**Supplementary Figure 1.**
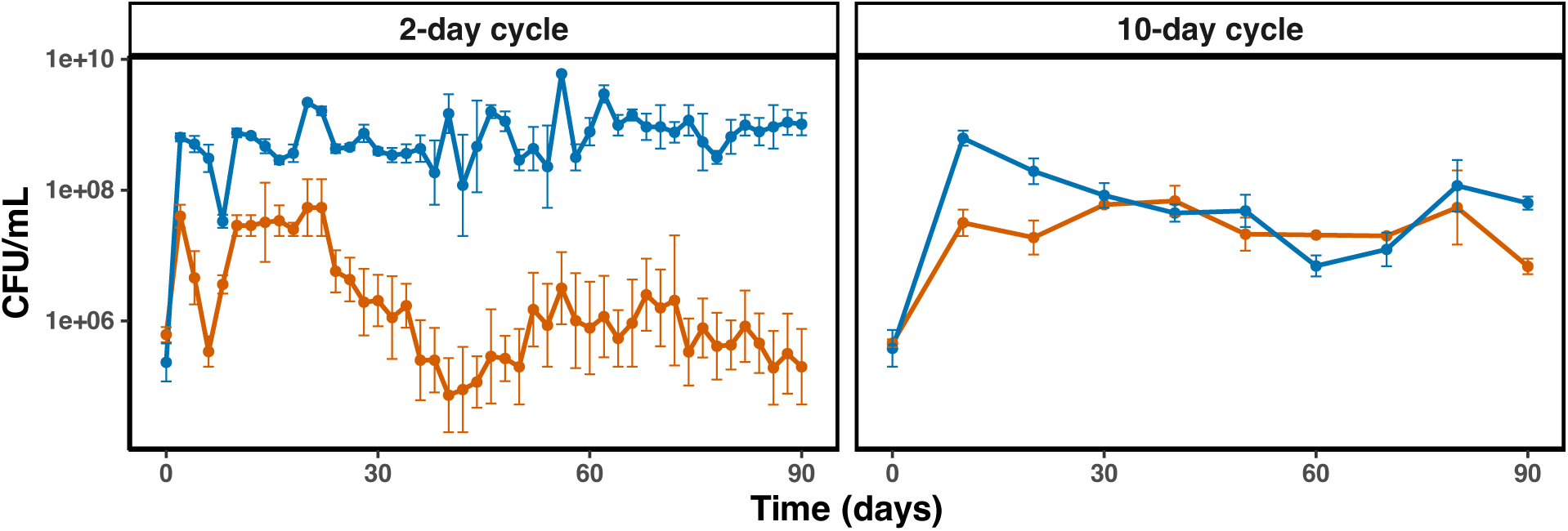
Cross-feeder cell density across 90 days of co-culture. Mean colony-forming units per milliliter (CFU/mL) of each ecotype across 90 days of co-culture in 2- and 10- day transfer regimes between overproducer Δ*metJ* (blue) and auxotroph Δ*metB* (orange) (n=3 per mutant). Error bars indicate (+/-) SEM

**Supplementary Figure 2.**
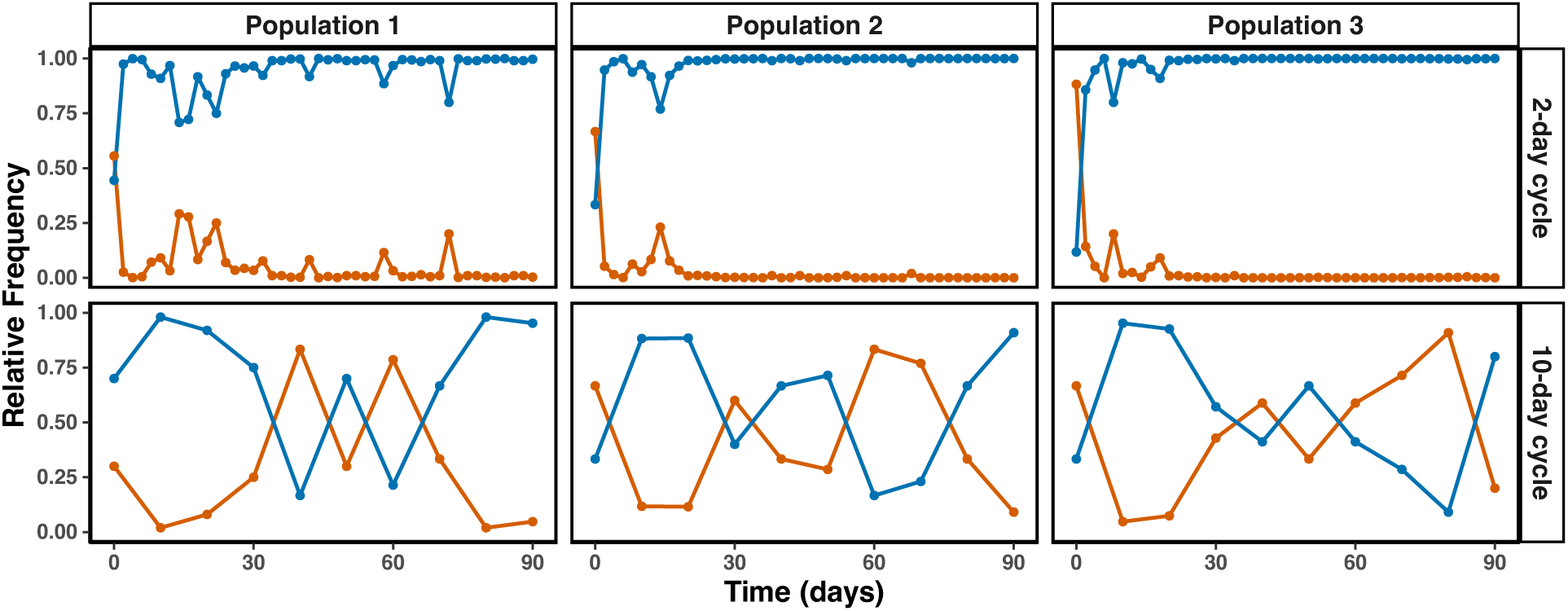
Cross-feeder relative frequency across 90 days of co-culture. Relative frequency of overproducer Δ*metJ* (blue) and auxotroph Δ*metB* (orange) for populations 1, 2, and 3 in co-culture conditions.

**Supplementary Figure 3.**
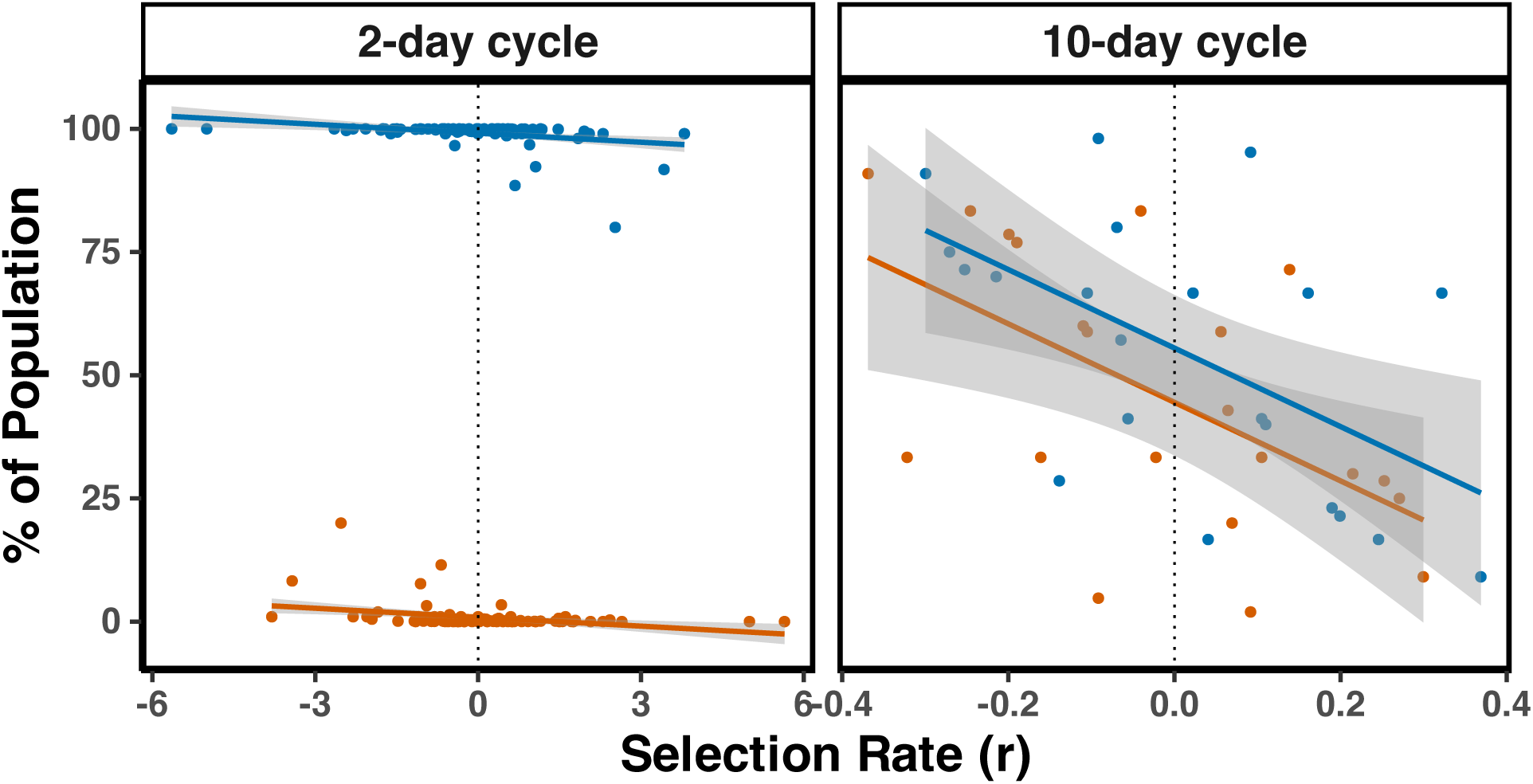
Selection rate versus subpopulation size across final 60 days of co-culture. Selection rate (*r*) of each the overproducer Δ*metJ* (blue) or auxotroph Δ*metB* (orange) at each mutant’s percentage of the population throughout the co-culture following pattern manifestation. Gray shaded regions indicate 95% confidence intervals. Selection rates were calculated by calculating the difference between the Malthusian parameters of each ecotype between each sampling point.

**Supplementary Figure 4.**
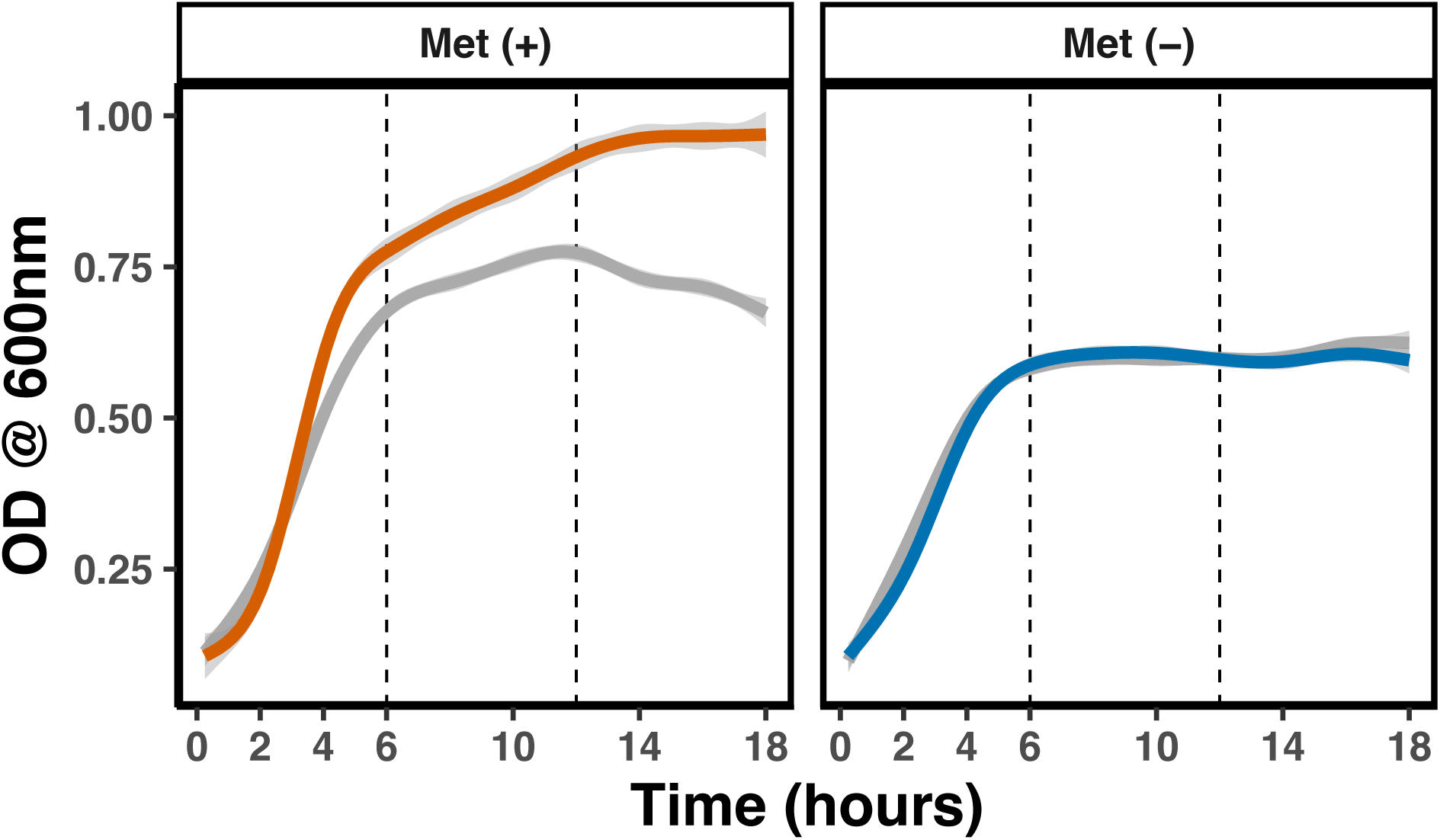
Mutant growth curves following a 1:10 dilution in supplemented (Met(+)) or unsupplemented (Met (-)) mM9 media. Growth curves (measured by optical density at a wavelength of 600 nm) for auxotroph Δ*metB* (orange), overproducer Δ*metJ* (blue), and wild-type (gray) following a 1:10 dilution into fresh mM9 minimal media following 20 hours of growth. Gray shading indicates 95% confidence intervals.

**Supplementary Figure 5.**
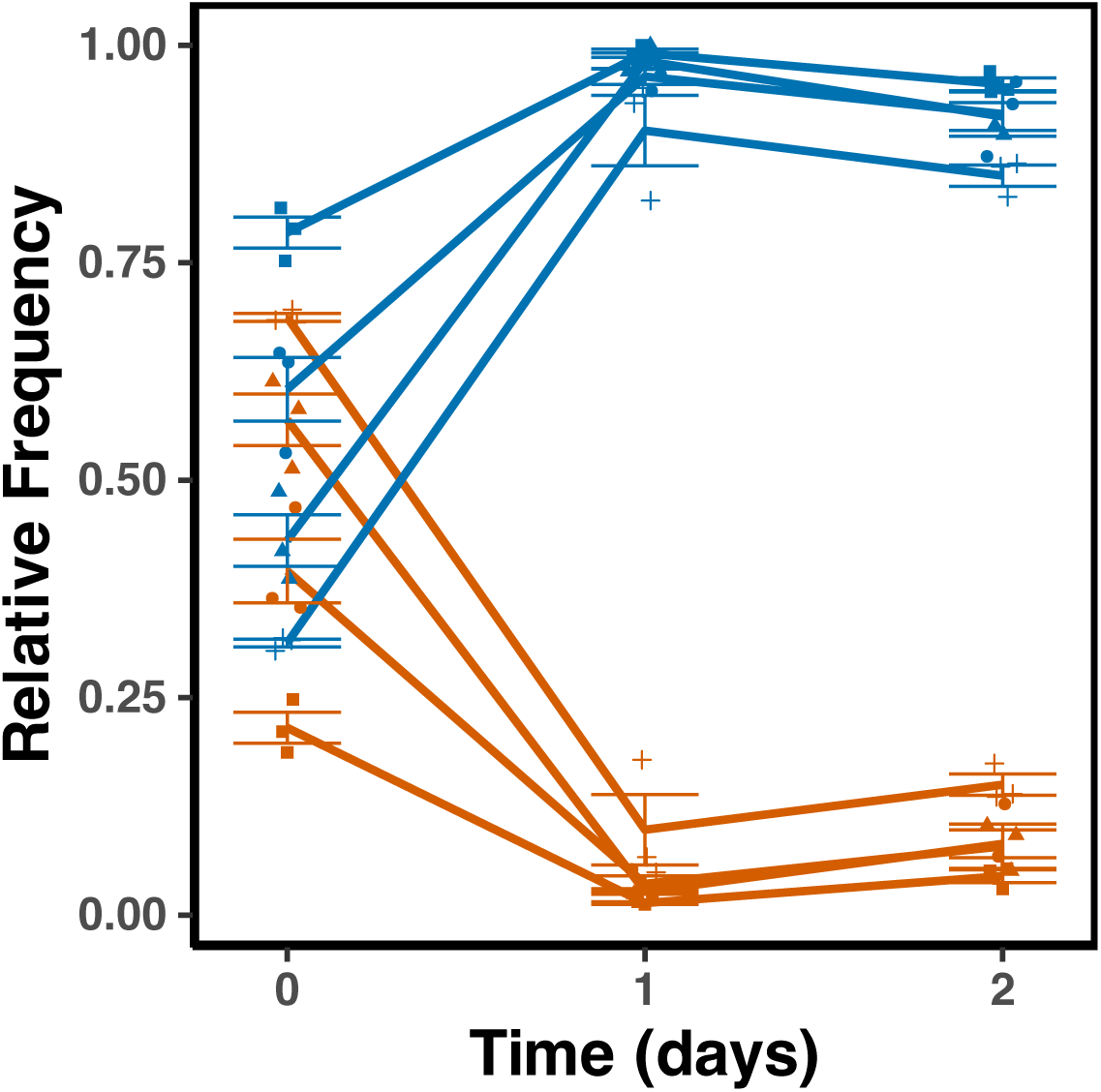
Ecotype frequencies across 2-days. Mean relative frequency of each ecotype across 2 days of co-culture across a range of starting mutant frequency (n=3 per starting frequency). Shapes indicate 3 replicates of each starting frequency. Error bars indicate (+/-) SEM.

**Supplementary Figure 6.**
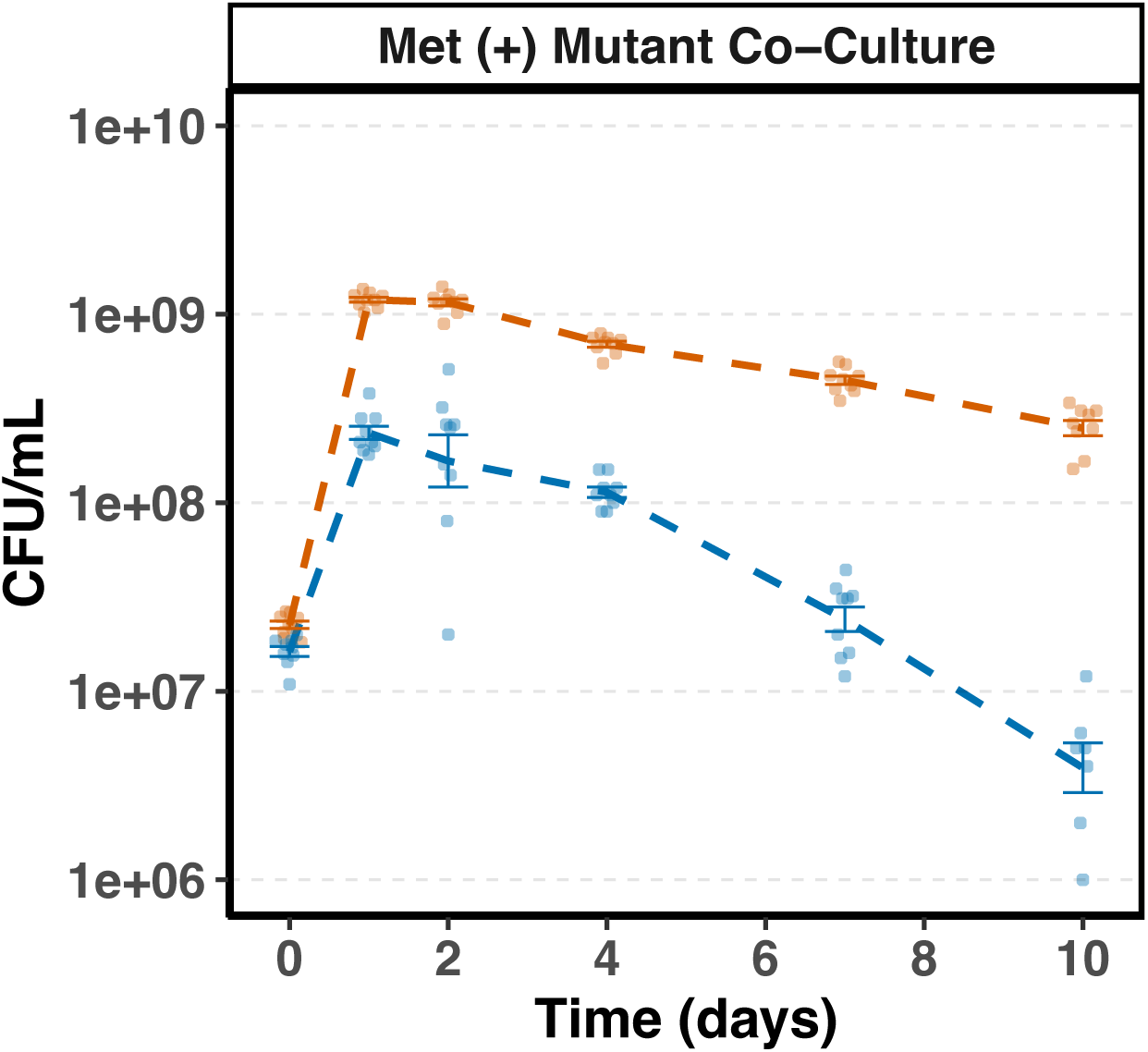
Mutant co-culture in methionine supplemented mM9. CFU abundance of clones (n=6) of overproducer Δ*metJ* or auxotroph Δ*metB* co-cultured directly across 10 days with an initial supplementation of 5.9 mM methionine into 10 mL mM9 + 0.2% glucose. Error bars indicate (+/-) SEM.

## Notes

https://github.com/BehringerLab/Ecotype_Coexistence

### Competing Interest Statement

The authors have declared no competing interest.

