## Supplemental_Figures for "Starvation drives co-existence in cross-feeding bacterial populations"

### Supplementary Figures

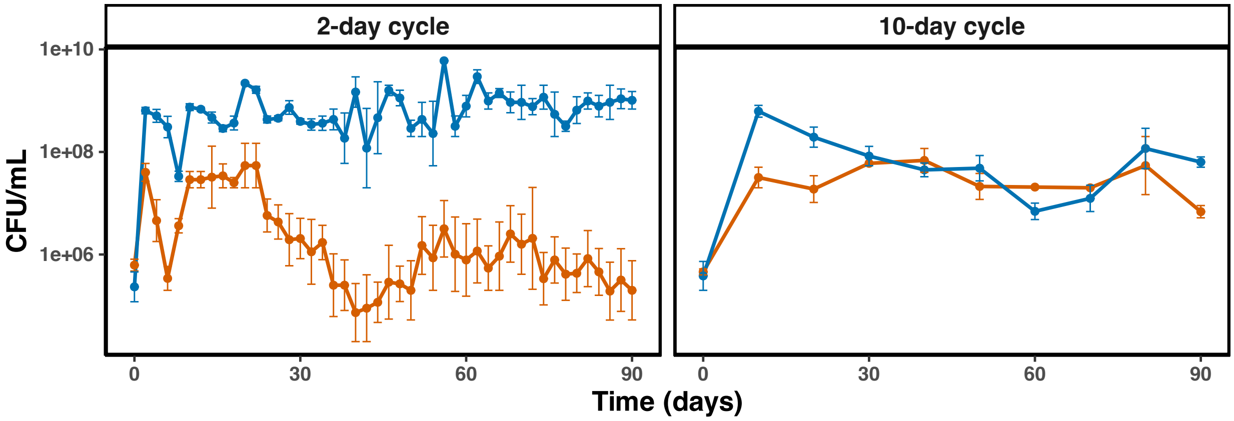


**Supplementary Figure 1. Cross-feeder cell density across 90 days of co-culture.** Mean colony-forming units per milliliter (CFU/mL) of each ecotype across 90 days of co-culture in 2- and 10-day transfer regimes between overproducer Δ*metJ* (blue) and auxotroph Δ*metB* (orange) (n=3 per mutant). Error bars indicate (+/-) SEM

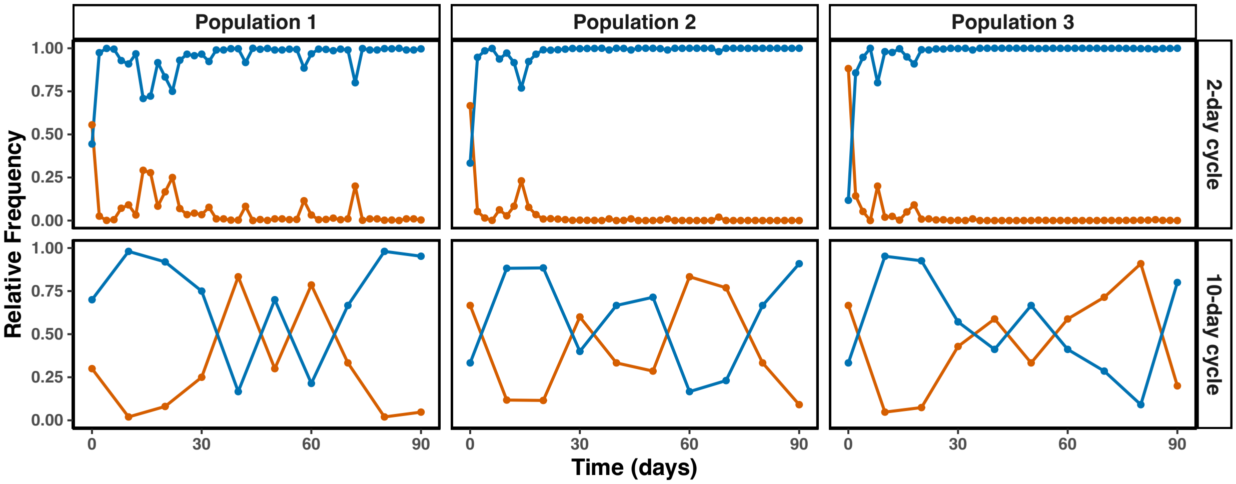


**Supplementary Figure 2. Cross-feeder relative frequency across 90 days of co-culture.** Relative frequency of overproducer Δ*metJ* (blue) and auxotroph Δ*metB* (orange) for populations 1, 2, and 3 in co-culture conditions.

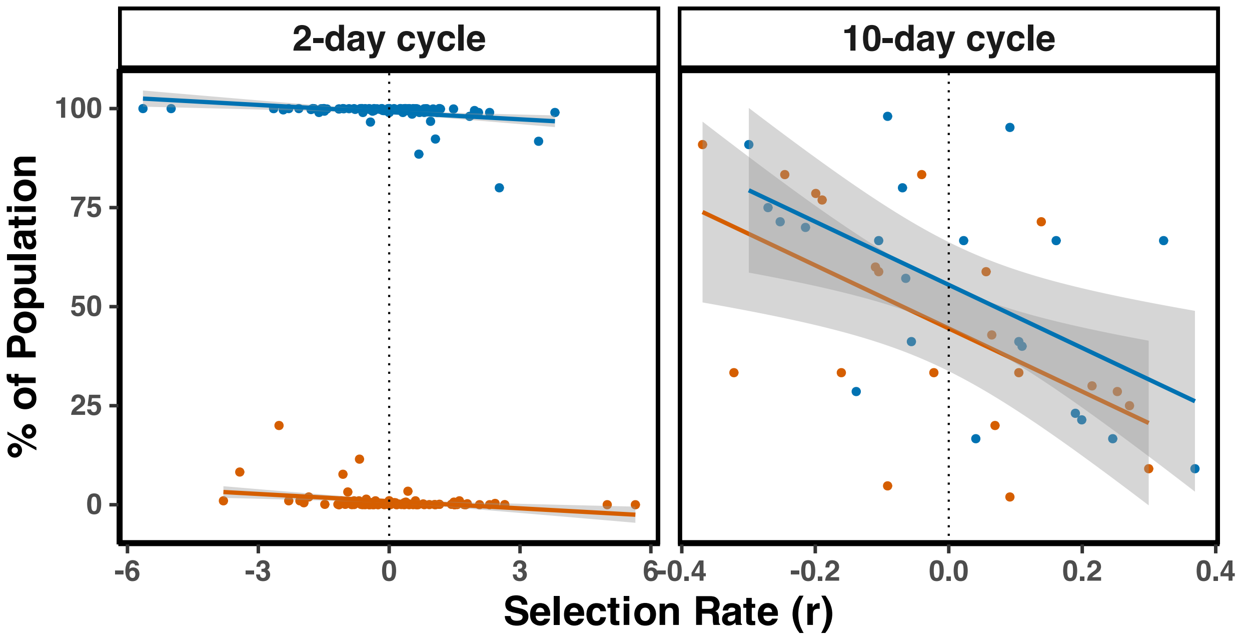


**Supplementary Figure 3. Selection rate versus subpopulation size across final 60 days of co-culture**. Selection rate (*r*) of each the overproducer Δ*metJ* (blue) or auxotroph Δ*metB* (orange) at each mutant's percentage of the population throughout the co-culture following pattern manifestation. Gray shaded regions indicate 95% confidence intervals. Selection rates were calculated by calculating the difference between the Malthusian parameters of each ecotype between each sampling point.

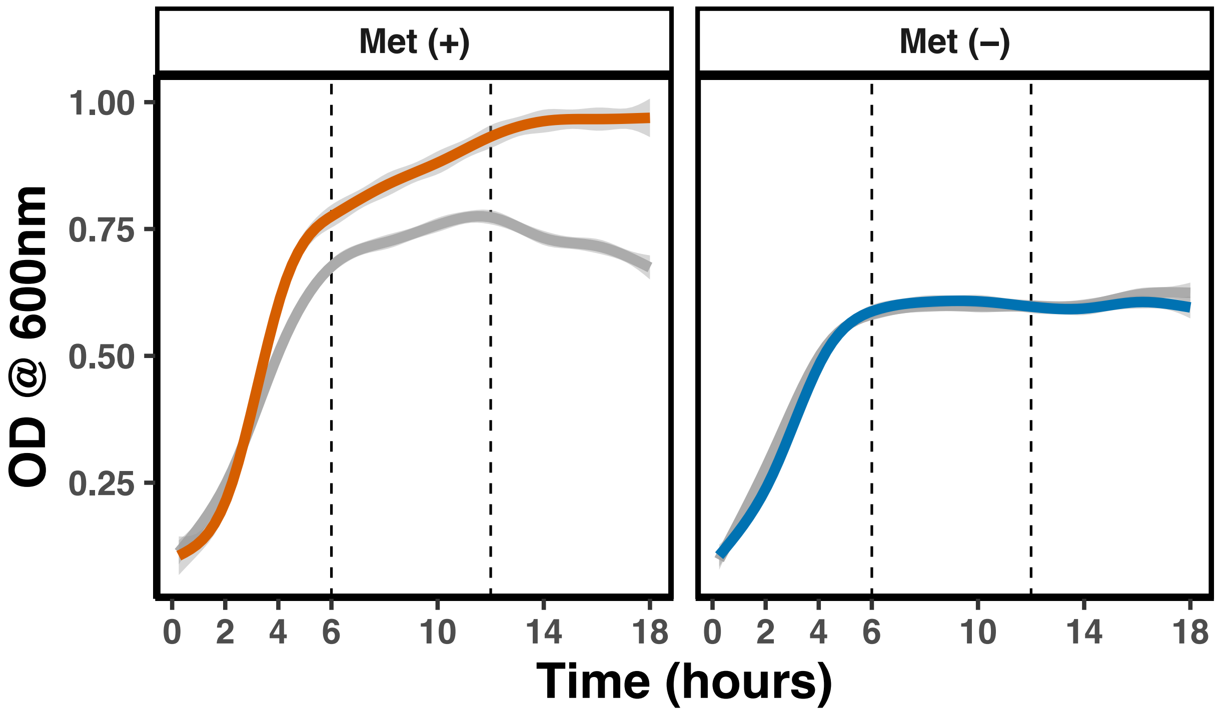


**Supplementary Figure 4. Mutant growth curves following a 1:10 dilution in supplemented (Met(+)) or unsupplemented (Met (-)) mM9 media.** Growth curves (measured by optical density at a wavelength of 600 nm) for auxotroph Δ*metB* (orange), overproducer Δ*metJ* (blue), and wild-type (gray) following a 1:10 dilution into fresh mM9 minimal media following 20 hours of growth. Gray shading indicates 95% confidence intervals.

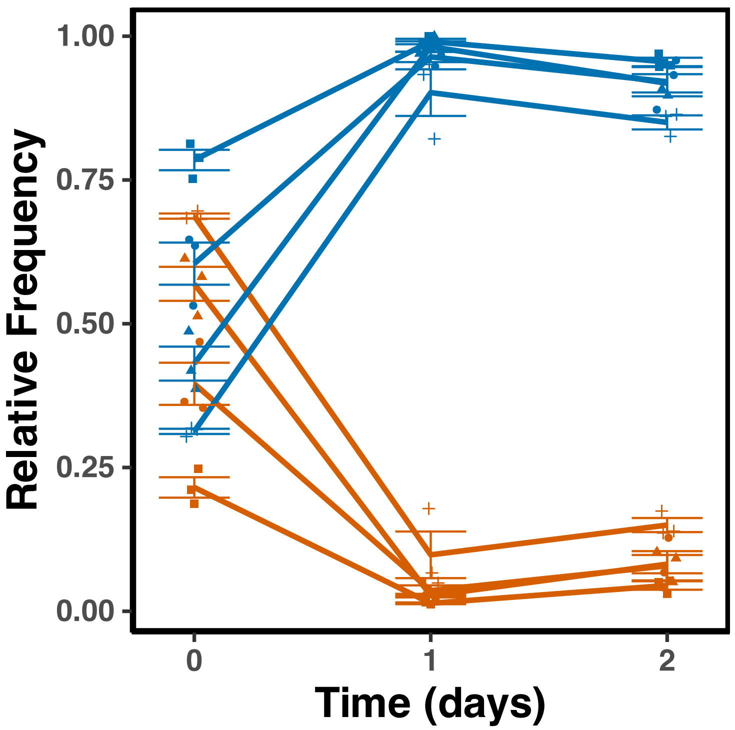


**Supplementary Figure 5. Ecotype frequencies across 2-days.** Mean relative frequency of each ecotype across 2 days of co-culture across a range of starting mutant frequency (n=3 per starting frequency). Shapes indicate 3 replicates of each starting frequency. Error bars indicate (+/-) SEM.

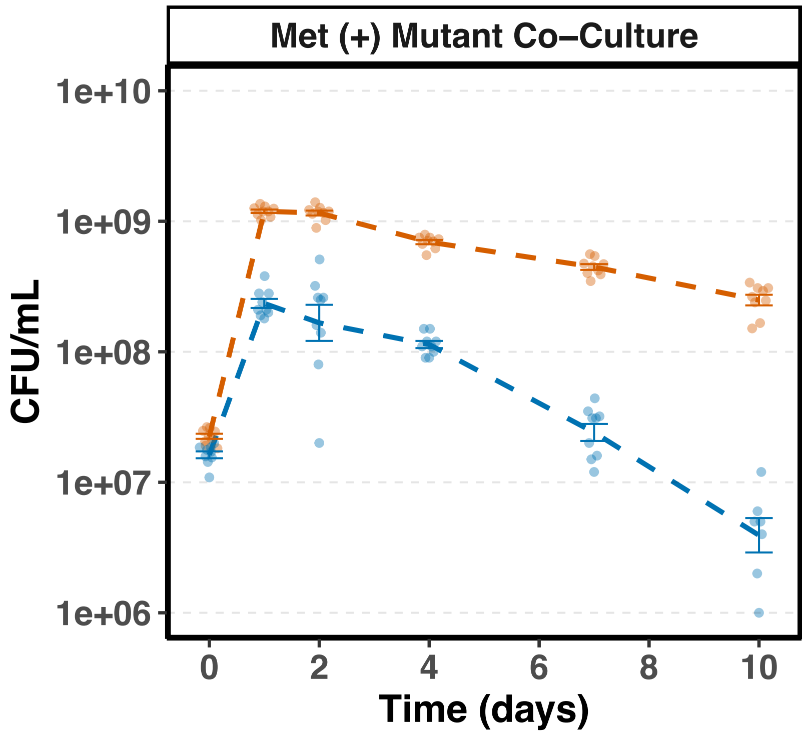


**Supplementary Figure 6. Mutant co-culture in methionine supplemented mM9.** CFU abundance of clones (n=6) of overproducer Δ*metJ* or auxotroph Δ*metB* co-cultured directly across 10 days with an initial supplementation of 5.9 mM methionine into 10 mL mM9 + 0.2% glucose. Error bars indicate (+/-) SEM.
